# Stromal asparagine supports tumor adaptation to oxidative phosphorylation inhibition through SLC38A4-mediated metabolic coupling

**DOI:** 10.64898/2026.03.18.710972

**Authors:** Ziyi Qin, Shengxi Li, Yeting Xu, Jiaming Zou, Jinyang Ma, Yingying Wang, Yucheng Wang, Rui Ju, Lin Wang, Lei Guo

**Affiliations:** Department of Pharmacology, Institute of Basic Medical Sciences, Chinese Academy of Medical Sciences & School of Basic Medicine, Peking Union Medical College, Beijing 100005, China; State Key Laboratory of Common Mechanism Research for Major Disease, Institute of Basic Medical Sciences, Chinese Academy of Medical Sciences and Peking Union Medical College, Beijing, China

**Keywords:** Pancreatic ductal adenocarcinoma, Metabolism, Carboxyamidotriazole orotate, Asparagine, SLC38A4, Cancer-associated fibroblasts, Integrated stress response, OXPHOS inhibition

## Abstract

**Purpose:** Pancreatic ductal adenocarcinoma (PDAC) is characterized by a nutrient-deprived and hypoxic tumor microenvironment (TME) that imposes severe metabolic stress on cancer cells. Under these conditions, tumor cells frequently activate the integrated stress response (ISR) to adapt to TME and develop resistance to therapies. However, how TME components support tumor adaptation to mitochondrial metabolic stress remains incompletely understood. Here, we aimed to identify key metabolite involved in ISR adaptation under oxidative phosphorylation (OXPHOS) inhibition and to elucidate the metabolic symbiosis between cancer-associated fibroblasts (CAFs) and PDAC cells.

**Methods:** We integrated transcriptomic and metabolomic analyses with functional assays. ISR activation was evaluated by assessing the phosphorylation of eIF2α (p-eIF2α) following treatment with carboxyamidotriazole orotate (CTO), an Complex I inhibitor. Metabolomic profiling was used to identify metabolites involved in ISR activation alleviation. Mouse models were used to assess therapeutic responses following depletion of the identified metabolite under CTO treatment. Genetic perturbation of *Slc38a4* was performed to assess its functional role in tumor cell adaptation to metabolic stress.

**Results:** We identified asparagine (ASN) as a critical metabolite supplied by CAFs to PDAC cells under OXPHOS inhibition. A minimum level of ASN is required for PDAC cells to execute ISR downstream adaptation. ASN depletion significantly enhanced the anti-tumor efficacy of OXPHOS inhibition both *in vitro* and *in vivo*. SLC38A4 emerged as a potential mediator of this interaction. SLC38A4 expression was associated with c-Myc, and its loss increased the sensitivity of PDAC cells to CTO-induced metabolic stress.

**Conclusion:** Our findings reveal a CAF-tumor metabolic crosstalk in which stromal-derived ASN supports PDAC cell adaptation to mitochondrial metabolic stress. Adaptive outcome of ISR signaling depends on the availability of key metabolic substrates such as ASN. When extracellular ASN supply is limited, the ATF4-dependent adaptive program collapses, converting ISR from a pro-survival response into a therapeutic vulnerability. SLC38A4 may function as a key mediator of this metabolic coupling and represents a potential target for enhancing the efficacy of OXPHOS inhibition in PDAC.

## Introduction

Pancreatic ductal adenocarcinoma (PDAC) is one of the most lethal cancers, largely due to late diagnosis, limited surgical eligibility, poor response to chemotherapy, and rapid development of therapeutic resistance. A main reason for this is its complicated and heterogeneous tumor microenvironment (TME), which plays a critical role in shaping tumor progression and dampens therapeutic outcomes(Kolbeinsson et al. 2023). Among the cellular components of the PDAC stroma, cancer-associated fibroblasts (CAFs) represent the most abundant population and serve as key regulators of tumor–stroma interactions(Mao et al. 2021). Beyond their structural role within the stroma, CAFs actively support tumor growth by reshaping the metabolic landscape of the TME. Increasing evidence suggests that CAFs function as metabolic support hubs for tumor cells by supplying diverse nutrients and metabolites and coordinating transporter expression between stromal and cancer cells. Through these mechanisms, CAFs form a dynamic metabolic symbiosis with PDAC cells that facilitates tumor survival and contributes to resistance against various therapeutic strategies, including conventional chemotherapy, metabolic inhibitors, and transporter-targeted therapies(Shan et al. 2017; Parker et al. 2020; Zhang et al. 2021; He et al. 2025). Consequently, disrupting the metabolic coupling between CAFs and tumor cells has emerged as a promising strategy to enhance anticancer therapies(Hu et al. 2024). However, the effectiveness of such approaches relies on a detailed understanding of the metabolic interaction networks between CAFs and PDAC cells, particularly under defined therapeutic pressures.

Oxidative phosphorylation (OXPHOS) has attracted considerable attention as a therapeutic target due to its central role in metabolic integration. PDAC cells often undergo metabolic rewiring in response to chemotherapy, such as gemcitabine, or targeted therapies including KRAS inhibitors, leading to an adaptive dependence on mitochondrial respiration. These observations have prompted increasing interest in pharmacological inhibition of OXPHOS as a therapeutic strategy. However, inter- and intratumoral heterogeneity in PDAC suggests that not all cancer cells respond uniformly to metabolic inhibitors (Masoud et al. 2020). In addition, the TME, particularly metabolic coupling between CAFs and PDAC cells, can attenuate the efficacy of metabolic therapies. Therefore, understanding how CAF-PDAC metabolic interactions evolve under OXPHOS inhibition may provide important insights for developing combination strategies and identifying patients who are most likely to benefit from OXPHOS-targeted therapies. In the present study, we used carboxyamidotriazole orotate (CTO), a previously characterized mitochondrial Complex I inhibitor with potent antiproliferative effects on tumor cells (Ju 2016; Shi 2019), to investigate tumor metabolic adaptation under OXPHOS inhibition.

Integrated stress response (ISR) is one of the mechanisms to adapt to environmental challenges and pathological conditions(Nakamura et al. 2010). ISR is a conserved signaling pathway that allows cells to respond to diverse environmental and pathological stresses, including nutrient deprivation, hypoxia, endoplasmic reticulum stress, and mitochondrial dysfunction(Malnassy et al. 2024). Central to ISR activation is the phosphorylation of the alpha subunit of eukaryotic initiation factor 2 (eIF2α), which suppresses global protein translation while selectively inducing stress-responsive transcription factors such as ATF4. Among the upstream kinases regulating ISR, general control nonderepressible 2 (GCN2) acts as a key sensor of amino acid deprivation. When intracellular amino acid levels decrease, uncharged tRNAs bind to GCN2 and trigger its activation, thereby initiating the GCN2-eIF2α-ATF4 signaling cascade that promotes amino acid biosynthesis, transport, and metabolic adaptation(Dong et al. 2000).

Although numerous studies have demonstrated that amino acid deficiency can activate the GCN2-ISR pathway, it remains unclear which specific amino acids play dominant roles in mediating ISR activation under defined metabolic stresses such as OXPHOS inhibition(Missiaen et al. 2022; Solvay et al. 2023; Novak et al. 2025). Furthermore, the extent to which extracellular amino acid supply from the TME modulates ISR signaling remains poorly understood. These observations raise the possibility that amino acid availability within the TME may critically determine the adaptive outcome of ISR signaling during therapeutic metabolic stress.

SLC38A4 is a sodium-coupled neutral amino acid transporter that mediates the uptake of asparagine (ASN), glutamine, and other neutral amino acids, thereby contributing to intracellular amino acid homeostasis. Although its physiological functions have been mainly studied in development and metabolic regulation, emerging evidence suggests that SLC38A4 may also participate in tumor metabolic regulation. Altered SLC38A4 expression has been associated with mTOR activation and tumor proliferation in colorectal and hepatocellular carcinoma, while elevated SLC38A4 expression has been linked to favorable prognosis in colorectal cancer liver metastasis(Li et al. 2021; Khosroshahi et al. 2024; Liu et al. 2025; Matoba et al. 2019; Xie et al. 2022). These findings suggest that SLC38A4 may influence tumor progression through its role in metabolite transport and that its function is likely shaped by the metabolic context of the TME. However, the role of SLC38A4 in PDAC remains largely unexplored.

In this study, we investigated metabolic interactions between CAFs and PDAC cells under CTO-induced OXPHOS inhibition. By integrating metabolomic and functional analyses, we identified ASN as a key metabolite involved in tumor adaptation to metabolic stress and identified SLC38A4 as a critical mediator of tumor-stromal metabolic coupling. Our findings provide new insights into how amino acid and its relatant transporter contributes ISR-dependent metabolic adaptation in PDAC and suggest that targeting SLC38A4 may represent a potential strategy for enhancing the efficacy of OXPHOS-based metabolic therapies.

## Methods

### Cell cultures and reagents

Mouse cell lines include KPC cell lines, Pan02 cell lines (Cell Resource Center, Peking Union Medical College [PCRC], Beijing, China), pancreatic stellate cell lines(IM-M088, Immocell, Xiamen, China), and BMDM (Extracted from bone marrow of C57BL/6 mice) were cultured in the complete cell DMEM media. The complete cell DMEM media were supplemented with 10% fetal bovine serum (FBS) (Yeasen, 40131ES), 1% (v/v) penicillin-streptomycin (Yeasen, 60162ES76), and 1% (v/v) glutamine (Solarbio, G0200). The cells were cultured at 37°C in a humidified environment with 5% CO2 in air. Cancer-associated fibroblasts (CAFs) were harvested after a 48-hour co-culture with tumor cells of pancreatic stellate cells (PSCs) in a Transwell system. CTO (20nM) was purchased from MedChemExpress (L-651582). L-asparaginase (ASNase) (10nM, approximately 0.3-0.4 U/mL) was purchased from MedChemExpress (HY-P1923). L-asparagine (ASN) (20µM) was purchased from MedChemExpress (HY-N0667). MG-132 (10 µM) was purchased from MedChemExpress (HY-13259). Chloroquine (CQ) (40µM) was purchased from MedChemExpress (HY-W031727). Cells were routinely screened for mycoplasma contamination every two weeks and were confirmed to be mycoplasma-free.

### Transwell co-culture of CAFs and tumor cells

Tumor cells and PSC were seeded separately into the upper and lower chambers of a Transwell system at a 1:1 ratio. After allowing approximately 4 hours for cell attachment, the inserts containing the upper-chamber cells were transferred into the lower chambers. Cells were co-cultured for 48 hours. PSC after co-culture was then collected and used as CAFs.

### Preparation of CAF-Conditioned Medium (CM)

CAFs were seeded into 10-cm culture dishes and cultured until approximately 80% confluence. Cells were then maintained in DMEM for 48 hours to generate conditioned medium (CM). The supernatant was collected and filtered through a 0.22-µm filter.

For induction experiments, CM was mixed with fresh complete DMEM at a ratio of 1:4 (CM:fresh medium) and applied to tumor cells for 24 or 48 hours.

For drug-treated CAFs, cells at approximately 80% confluence were treated with CQ or CTO for 4 hours. The drug-containing medium was then replaced with drug-free DMEM, and cells were further cultured for 6 hours. The supernatant was subsequently collected and filtered through a 0.22-µm filter. The resulting CM was mixed with fresh complete DMEM at a ratio of 1:4 and applied to tumor cells for 48 hours.

For ultrafiltration experiments, CAF-CM was concentrated using centrifugal ultrafiltration tubes (ultrafiltration spin columns) (Henghuibio, UFC301096), and the filtrate was reconstituted with basal DMEM to the original volume.

For heat-treatment experiments, CAF-CM was heated at 98°C for 30 min using a water bath or metal heating block.

All CAF-CM samples were stored at −80°C and subjected to no more than three freeze–thaw cycles before use.

### Quantitative reverse transcription-polymerase chain reaction (RT-qPCR)

A total of 4×10^5^ cells were cultured onto 6 cm dishes for 48 h with or without CTO treatment. Total RNA was extracted from cells, reverse transcribed, and subjected to RT-qPCR analysis by using AG RNAex Pro Reagen, Evo M-MLV RT Premix, and SYBR® Green Premix Pro Taq HS qPCR Kit (Accurate Biology, AG21102, AG11706, and AG11701, respectively). The reaction temperature and number of cycles were set in accordance with the manufacturer’s instructions. *Slc38a4* expression levels were normalized to the *β-actin* mRNA level.

Primer sequences (RT–qPCR):

Slc38a4: F-AGCCGATTACGCGGATGAG; R-GGACAAGCCTAAGATCCCACTG

β-actin: F-GGCTGTATTCCCCTCCATCG; R-CCAGTTGGTAACAATGCCATGT

### Western blotting assay

A total of 4 × 105 cells were cultured in 6 cm culture dishes for 48 h with or without CTO treatment. After cell lysis using 200 µL RIPA lysis buffer (Solarbio, R0020) containing 1% (v/v) PMSF (Solarbio, P0100) and the protein phosphatase inhibitor (Solarbio, P1260), protein concentration was determined with the BCA Protein Quantification Kit (Yeasen, 20201ES86) for normalization. After protein quantification and denaturation using the 5×SDS-PAGE Protein Loading Buffer (Yeasen, 20315ES20), the proteins were separated on SDS-PAGE gels prepared with the BioSci® New Flash Protein AnyKD PAGE (DAKEWE, 8012011) and subsequently transferred onto a PVDF membrane (Millipore, IPVH00010). The membrane was then blocked with 5% skimmed milk (Yeasen, 36120ES76) prepared in TBST for 2 hours at room temperature with gentle shaking, followed by an overnight incubation at 4°C with the following diluted primary antibody(1:1000): eIF2α (Selleck, F3016), p-eIF2α(Ser51) (Selleck, F025), c-Myc (Abcam, AB32072), GCN2 (Abcam, AB302609), p-GCN2 (Bioss, bs-3155R), Slc38a4 (Proteintech, 20857-1-AP), β-actin(Cell Signaling Technology, 3700S). The membrane was then incubated with the appropriate secondary antibody at room temperature for 1 to 2 hours. Immunoreactive bands were visualized using an enhanced chemiluminescence kit (Tanon). The relative protein expression levels were quantified with ImageJ software and normalized to the corresponding β-actin levels.

### CUT&Tag and qPCR quantification

CUT&Tag was performed using a commercial CUT&Tag kit (Novoprotein, N259-YH01) according to the manufacturer’s instructions.For CUT&Tag-qPCR, enrichment at the Slc38a4 locus was assessed by quantitative PCR using three primer pairs spanning the target region(s). CUT&Tag-qPCR signals were presented as relative enrichment (normalized to IgG control, and expressed relative to a reference condition as indicated in the figure legends).

Primer sequences (CUT&Tag-qPCR):

Slc38a4-1: F-GTGGGGACAAGCGCAC; R-AGGTACAACCAACAAATTTTGACC Slc38a4-2: F-GCGCGCGTGTATGTGTG; R-TTCCGAGACCTGAATGCCCC Slc38a4-3: F-GTGTGCGCGCGTGTATGT; R-CCGAGACCTGAATGCCCCAC

### Seahorse OCR measurement

Oxygen consumption rate (OCR) was measured using the Seahorse XF24 extracellular flux analyzer (Agilent Technologies) according to the manufacturer’s instructions. Cells were seeded into Seahorse XF cell culture microplates at 8 × 10^3^ cells/well and incubated overnight to allow adherence.Cells were treated with CTO and/or CAF conditioned medium (CAF-CM) for 48 h prior to OCR measurement. Before the assay, culture medium was replaced with Seahorse XF assay medium supplemented with 10 mM glucose, 1 mM pyruvate, and 2 mM glutamine, and cells were equilibrated in a CO₂-free incubator for 45 min.

OCR was measured using the Seahorse XF Cell Mito Stress Test Kit (Alicelliagent, ALS22111). During the assay, the following compounds were sequentially injected: oligomycin, FCCP, rotenone/antimycin A. OCR values were normalized to total protein content.

### Sulforhodamine B (SRB) assay

Cell proliferation was assessed using a Sulforhodamine B (SRB) assay kit (Yeason, S7524100) according to the manufacturer’s instructions. Cells were seeded into 96-well plates at 8 × 10^3^ cells/well and incubated overnight to allow adherence.Cells were treated with CTO and/or CAF conditioned medium (CAF-CM) for 48 h. After the treatments, cells were fixed with trichloroacetic acid (TCA) and stained with SRB solution. Excess dye was removed by washing with 1% acetic acid, and the protein-bound dye was subsequently dissolved in Tris base solution. Absorbance was measured at 515 nm using a microplate reader. Cell proliferation was calculated relative to the control group.

### NAD^+^/NADH ratio assay

Intracellular NAD⁺/NADH ratios were determined using commercial NAD⁺/NADH Assay Kit (Bioss, Beijing, China) according to the manufacturer’s instructions. Briefly, cells were harvested and lysed to obtain total cellular extracts. To measure NADH, an aliquot of each lysate was subjected to selective decomposition of NAD⁺ as instructed by the kit, whereas a parallel aliquot was used for total NAD (NAD⁺ + NADH) determination. NAD⁺ levels were calculated by subtracting NADH from total NAD. The NAD⁺/NADH ratio was calculated accordingly. Values were normalized to total protein content.

### Xenograft and orthotopic pancreatic cancer models

All animal experiments were approved by the Institutional Animal Care and Use Committee (IACUC) of the Institute of Basic Medical Sciences, Chinese Academy of Medical Sciences(approval number: ACUC-XMSB-2025-162), and all procedures were performed in accordance with institutional guidelines for animal care and use. Mice were housed in the specific pathogen-free (SPF) facility of the Experimental Animal Centre, School of Basic Medicine, Peking Union Medical College. Male C57BL/6J mice (6–8 weeks old) were used for tumor modeling. For orthotopic implantation, 1×10⁶ KPC cells suspended in 25 µl of matrix mixture were injected into the pancreatic tail parenchyma. Following a 3-day postoperative recovery period, mice were randomly assigned to treatment groups. Mice received carboxyamidotriazole orotate (CTO, 90 mg/kg) or an equal volume of polyethylene glycol 300 by oral gavage once daily. In addition, mice were administered Oncaspar (PEG-L-asparaginase, MCE, HY-108860, 2 U/g) or an equal volume of physiological saline by intraperitoneal injection every 7 days. Treatment lasted for 10 days, and mice were monitored daily for health status. At the experimental endpoint (13 days after tumor implantation), mice were sacrificed and tumors were excised, weighed, and photographed for further analysis.

### Immunohistochemical (IHC) staining and Masson trichrome staining

Tissue sections were prepared from fixed animal tumor tissues through a standard process of dehydration, permeabilization, and paraffin embedding. Immunohistochemical staining was performed on tissue sections using primary antibodies against Ki67 (GB111499, diluted 1:500). Masson’s trichrome staining was also conducted to evaluate collagen deposition. All stained sections were scanned using CaseViewer software and quantitatively analyzed with ImageJ software.

### Survival Analysis Using UCSC Database

Overall survival (OS) analysis of EIF2S1 (eIF2α) (in pancreatic ductal adenocarcinoma (PDAC) patients was performed using the UCSC online platform (http://gepia2.cancer-pku.cn). Gene expression data were derived from GDC PAAD datasets. Patients were stratified into high-, median- and low-expression groups based on the median expression level ofEIF2S1. Kaplan-Meier survival curves were generated, and statistical significance was evaluated using the log-rank test.

### Transcriptome Sequencing and Bioinformatics Analysis

RNA sequencing was performed by Novogene Co., Ltd. Raw sequencing reads were aligned to the mouse reference genome(mm10)using HISAT2(v2.0.5). Gene expression quantification was conducted using featureCounts(v1.5.0-p3). Differential expression analysis was performed using DESeq2(v1.20.0). Genes with |log_2_fold change)|>1 and adjusted p-value(padj)<0.05 were considered significantly differentially expressed.

### Motif prediction and primer design

Putative c-Myc binding motifs within the Slc38a4 regulatory region were predicted using the JASPAR database. Genomic sequences surrounding the Slc38a4 transcription start site (TSS) (e.g., ±2 kb; or specify the exact window used) were scanned for MYC/MYC::MAX (E-box) motifs. Predicted sites with a relative score ≥ 0.85 were considered candidate binding regions and were used to guide the design of CUT&Tag-qPCR primer sets targeting the Slc38a4 locus.

### Statistical analysis

All data are presented as mean ±SD. Statistical analyses were performed using GraphPad Prism. Comparisons between two groups were conducted using a two-tailed unpaired Student’s t-test. Comparisons among multiple groups were conducted using one-way ANOVA followed by Tukey’s multiple comparisons test. A p-value < 0.05 was considered statistically significant. Sample sizes (n) are indicated in the figure legends and represent biological replicates. For animal studies, sample sizes were determined based on preliminary experiments and previous studies, and no statistical method was used to predetermine sample size. Animals were randomly assigned to treatment groups, and investigators were not blinded to group allocation during experiments and outcome assessment unless otherwise stated.

### Untargeted metabolomics analysis

For intracellular metabolomics analysis, PDAC cells were seeded in 6 cm dishes and treated with the indicated conditions (Control, CTO, CTO+ASNase) for 48 h. Approximately 4×10^5^ cells per sample (n = 5-6) were collected for metabolite extraction. The culture dishes were rapidly placed on dry ice, and metabolites were extracted directly by adding 40µL of precooled 80% (v/v) methanol per 1×10^6^ cells. Cells were scraped using a cell spatula and transferred to microcentrifuge tubes. The extracts were centrifuged twice at 15,000 rpm for 30 min at 4℃ to remove cellular debris. Subsequently, 50 µL of the supernatant was transferred into LC-MS injection vials for untargeted metabolomic analysis using high-performance liquid chromatography-tandem mass spectrometry (HPLC-MS/MS).

For metabolomic profiling of CAF-derived metabolites, CAF conditioned medium (CAF CM) was collected from CAF cultures under the indicated treatment conditions (Control, CTO, CQ, and CTO + CQ; n = 5-6), with basic DMEM serving as a medium control (n = 2). CAFs were treated with the indicated drugs for 24 h, followed by replacement with fresh basic DMEM and further incubation for 6 h to collect conditioned medium. The culture supernatant was centrifuged at 3,000 rpm for 10 min to remove cellular debris and filtered through a 0.22µm membrane. Metabolites in the supernatant were extracted by adding four volumes of precooled 80% (v/v) methanol, followed by vortexing and incubation on ice. The samples were centrifuged twice at 15,000 rpm for 30 min at 4°C. The resulting supernatant was transferred to LC-MS injection vials for subsequent untargeted metabolomics analysis using HPLC-MS/MS.

### Human sample

All formalin-fixed paraffin-embedded (FFPE) patient tissue samples were obtained from the with pancreatic cancer by PUMCH were eligible for the study. Written informed consent was obtained from all patients. The study was approved by the PUMCH Institutional Review Board (approval number: I-24PJ2172). FFPE tumor specimens were used for subsequent immunofluorescence staining.

## Results

### 1. Tumor cells resist CTO-induced proliferation inhibition through activation of the ISR pathway

In our preliminary experiments, we observed that CTO significantly inhibited the proliferation of tumor cells. As CTO has previously been identified as an inhibitor of oxidative phosphorylation (OXPHOS), we performed intracellular metabolomic profiling to investigate metabolic alterations in cancer cells following CTO treatment (Fig. 1A). We examined amino acids associated with the GCN2 signaling pathway and found that asparagine (ASN) and aspartate were markedly reduced after CTO treatment (|log₂FC| > 1). These findings suggest that CTO-induced metabolic stress may lead to amino acid deprivation, potentially resulting in the accumulation of uncharged tRNAs and subsequent activate the GCN2 pathway. Based on this observation, we hypothesized that OXPHOS inhibition by CTO could trigger activation of the integrated stress response (ISR). To test this hypothesis, we assessed the expression levels of phosphorylated eIF2α (p-eIF2α), a key indicator of ISR activation, and ATF4, a downstream transcription factor induced during ISR signaling, using Western blot analysis (Fig. 1B-C). The results showed that both p-eIF2α and ATF4 were significantly upregulated following CTO treatment, indicating activation of ISR signaling under CTO-induced metabolic stress.

**Figure 1.**
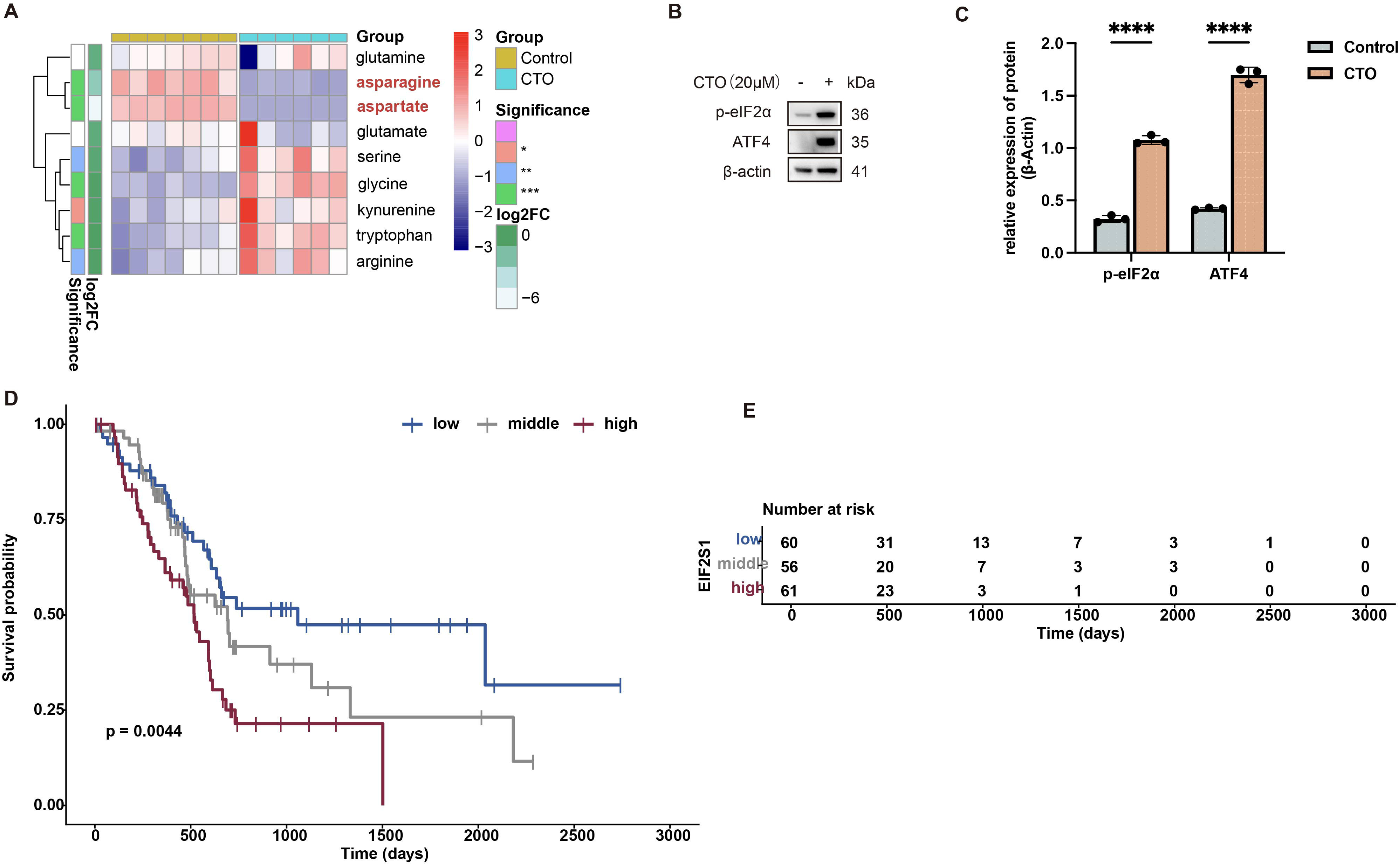
CTO induces metabolic stress and activates the ISR pathway in pancreatic cancer cells. A. Intracellular metabolomic analysis of GCN2 pathway-related metabolites in Pan02 cells (t test, ****p < 0.0001, ***p < 0.001, **p < 0.01, *p < 0.05). B. Representative Western blot images of ISR-related proteins in the KPC cell line following CTO treatment. C. Quantification of Western blot results shown in panel B using ImageJ (t test, n = 3, ****p < 0.0001). D. Kaplan-Meier survival analysis showing the association between EIF2S1 expression levels and overall survival of pancreatic cancer patients based on UCSC database data. E. Statistical summary of survival data corresponding to panel D.

To further to explore its association with patient prognosis, we analyzed pancreatic cancer data from the TCGA database. Patients were stratified according to EIF2S1 expression levels to explore the clinical relevance of ISR signaling. Kaplan-Meier survival analysis revealed that high EIF2S1 expression was significantly associated with poorer overall survival in pancreatic cancer patients (Fig. 1D-E).

Previous studies have shown that ATF4 activation promotes the expression of a series of genes that support tumor cell survival under unfavorable microenvironmental conditions, particularly asparagine synthetase (ASNS), which contributes to maintaining intracellular ASN homeostasis. Taken together, these results suggest that activation of the ISR pathway represents an adaptive survival strategy employed by tumor cells to cope with metabolic stress. Under CTO-induced stress, tumor cells activate ISR signaling to adapt to unfavorable conditions.

### 2. CAFs alleviate metabolic stress in tumor cells by supporting cancer cells with metabolites

To further simulate tumor-stromal interactions *in vitro*, we collected CAF conditioned medium (CM) and performed SRB proliferation assays on cancer cells under CTO-induced inhibition (Fig. 2A). We observed that CAF CM significantly rescued the inhibitory effect of CTO on tumor cell proliferation. To determine whether CAF CM interferes directly with the inhibitory effect of CTO on mitochondrial respiration, we performed Seahorse OCR analysis(Fig.2B-E). Tumor cells were divided into three groups: untreated cells, cells treated with CTO for 48 hours, and cells treated with CTO together with CAF CM for 48 hours. As expected, CTO significantly suppressed Complex I activity. The presence of CAF CM did not significantly restore mitochondrial Complex I activity compared with CTO treatment alone. These results indicate that CAF CM does not rescue tumor cell proliferation by directly antagonizing CTO activity or its function. We further examined intracellular redox status by measuring the NAD^+^/NADH ratio. CTO treatment significantly reduced the NAD^+^/NADH ratio, indicating relative depletion of NAD^+^(Fig. 2F-G). In contrast, CAF CM significantly increased the NAD^+^/NADH ratio compared with CTO treatment alone. These findings suggest that CAF CM restores intracellular redox balance independently of mitochondrial Complex I activity.

**Figure 2.**
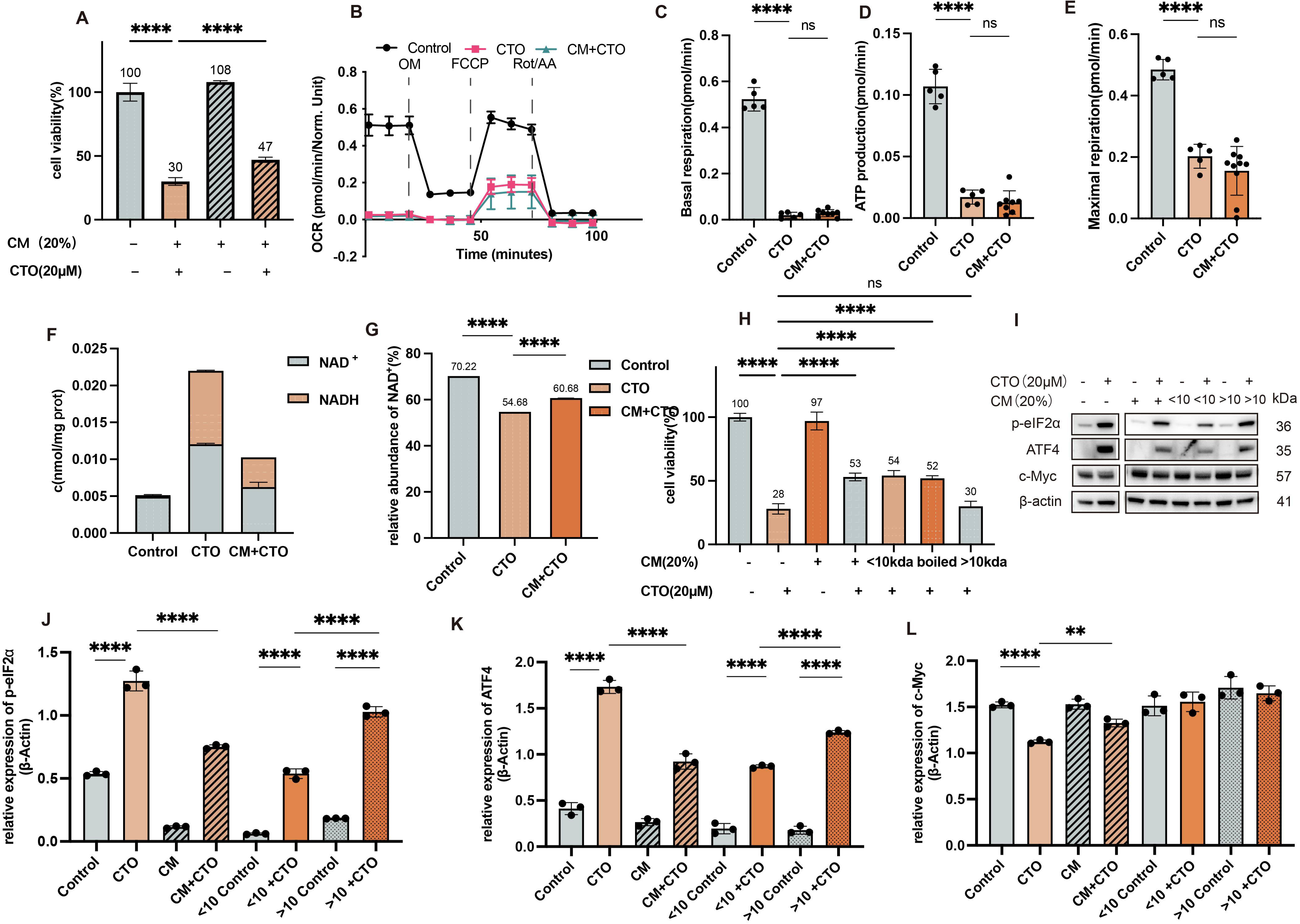
CAF conditioned medium partially rescues CTO-induced mitochondrial dysfunction. A. SRB cell proliferation assay of KPC cells following CTO treatment (t test, n = 3, ****p < 0.0001). B. Seahorse XF analysis showing mitochondrial respiration profiles of Pan02 cells. C. Quantification of basal respiration. D. Quantification of ATP production rate. E. Quantification of maximal respiration. F. Intracellular NAD⁺ + NADH levels in Pan02 cells. G. Relative NAD⁺/NADH ratio in Pan02 cells. H. SRB proliferation assay of KPC cells under the indicated treatment conditions (t test, n = 3, ****p < 0.0001, ***p < 0.001). I. Representative Western blot images of ISR-related proteins in Pan02 cells under the indicated treatment conditions. J-K. Quantification of Western blot results shown in panel J (t test, n = 3, ****p < 0.0001).

To identify the key component responsible for this rescue effect, CAF CM was subjected to boiling and ultrafiltration to separate macromolecules and small metabolites. Ultrafiltration generated two fractions: a <10 kDa fraction containing mainly low-molecular-weight peptides and metabolites, and a >10 kDa fraction enriched for larger proteins. Boiled CAF CM contained predominantly metabolites. SRB assays revealed that both the <10 kDa fraction and boiled CAF CM retained the ability to rescue CTO-induced growth inhibition, whereas the >10 kDa fraction failed to do so (Fig. 2H). Western blot analysis further showed that small metabolites in CAF CM significantly attenuated ISR activation under CTO stress and increased c-Myc expression(Fig. 2I-L). c-Myc inhibition was previously suggested to be one of the key reasons of CTO-induced tumor cell proliferation inhibition, and also reported to be a consequence of global translation inhibition under ISR activation. Interestingly, although the >10 kDa fraction also increased c-Myc expression, which highly related to malignant proliferation of tumor, it failed to restore tumor cell proliferation under CTO treatment, suggesting that c-Myc upregulation alone is insufficient to overcome CTO-induced metabolic stress.

These results suggest that metabolite supplementation provided by CAFs is a key factor that weakens the inhibitory effect of CTO on tumor cell growth through extracellular metabolic support. Notably, although ISR signaling was strongly activated under CTO treatment, tumor cells still exhibited impaired proliferation, suggesting that endogenous metabolic adaptation driven by ATF4 alone is insufficient to fully compensate for CTO-induced metabolic stress. In contrast, metabolite supply from CAF-conditioned medium effectively restored tumor cell proliferation, highlighting the importance of stromal metabolic support.

### 3. CAF-derived ASN alleviates ISR stress and supports tumor cell proliferation under OXPHOS inhibition

Previous studies have reported that CAFs can provide metabolic support to tumor cells through autophagy-mediated metabolite release. Therefore, we hypothesized that the key metabolites present in CAF CM might originate from CAF autophagy. Meanwhile, since CTO is an OXPHOS inhibitor, its effects on CAF biology may also influence the metabolic composition of CAF CM. To investigate this possibility, CAFs were treated with the autophagy inhibitor chloroquine (CQ) or CTO for 24 hours, followed by replacement with drug-free DMEM and subsequent collection of CAF CM (Fig. 3B). We then examined the effects of these CAF CM on tumor cells under CTO-induced metabolic stress. We found that CM derived from CQ-treated CAFs still retained the ability to rescue CTO-induced growth inhibition, whereas CM derived from CTO-treated CAFs lost this rescue effect(Fig. 3A). These findings suggest that the metabolite responsible for metabolic compensation is unlikely to be generated through CAF autophagy.

**Figure 3.**
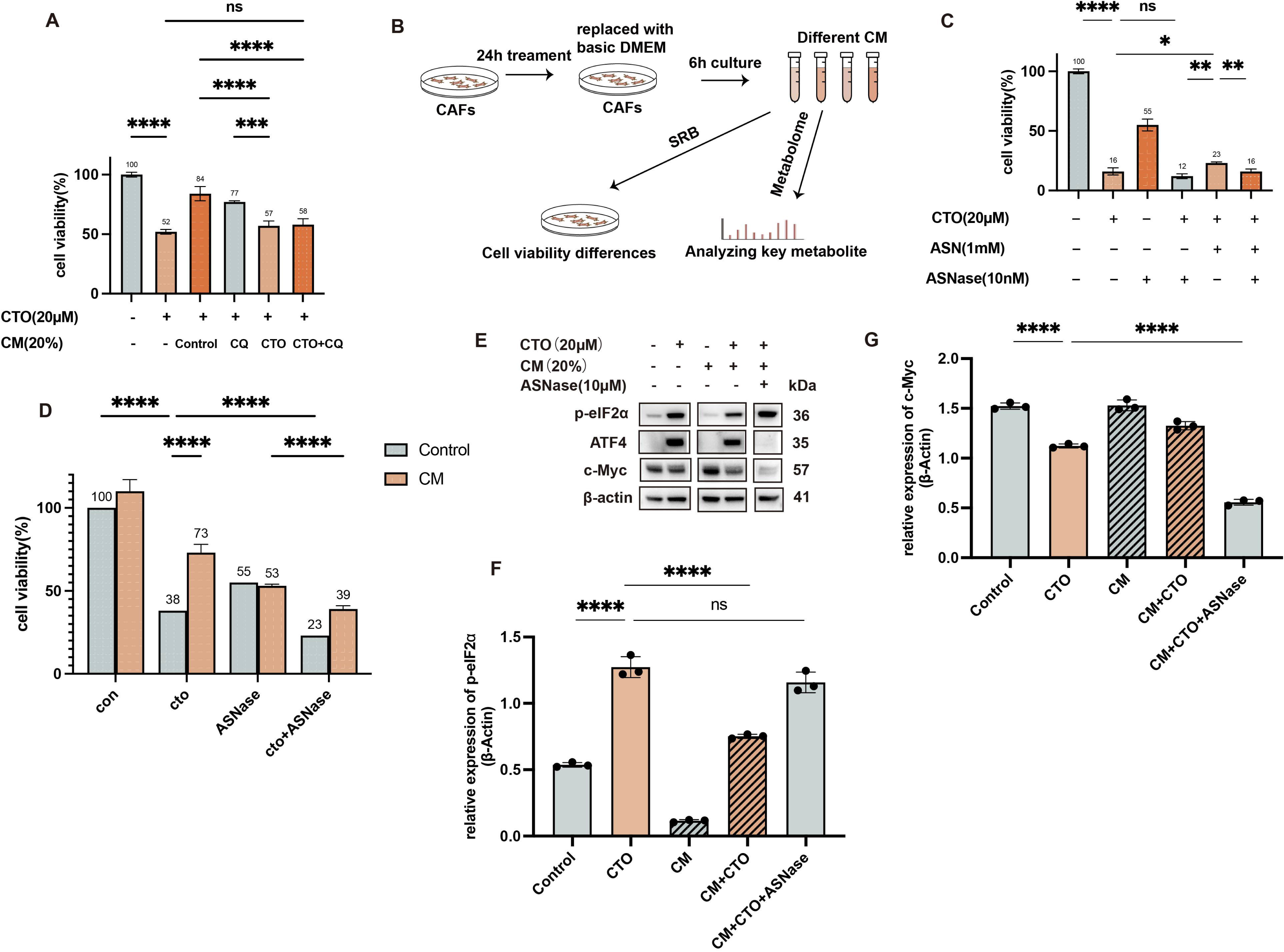
CAF-derived metabolites alleviate CTO-induced metabolic stress. A. SRB proliferation assay of Pan02 cells under the indicated conditions (t test, n = 3, ****p < 0.0001). B. Schematic diagram showing the workflow for collecting secreted metabolites from CAF conditioned medium. C. SRB proliferation assay of Pan02 cells treated with specific metabolites identified from CAF conditioned medium (t test, n = 3, ****p < 0.0001, **p < 0.01, *p < 0.05). D. SRB proliferation assay of Pan02 cells under the indicated conditions (t test, n = 3, ****p < 0.0001). E. Representative Western blot images of Pan02 cells. F–G. Quantification of Western blot results shown in panel E using ImageJ (t test, n = 3, ****p < 0.0001).

Based on these observations, metabolomic profiling was performed on CAF CM under different treatment conditions. A filtering strategy was applied to identify candidate metabolites that were enriched in untreated CAF CM and CQ-treated CAF CM relative to DMEM but decreased in CTO-treated CAF CM(Fig. 3B). Using this strategy, 4 metabolites fully met the screening criteria and 11 additional metabolites partially met the criteria (Table. S1). Among these candidates, ASN was identified as the most likely key metabolite based on literature evidence. To validate this hypothesis, tumor cells were subjected to ASN supplementation and depletion experiments under CTO stress (Fig. 3C). ASN supplementation significantly rescued CTO-induced growth inhibition, whereas degradation of ASN by ASNase abolished this rescue effect. We next examined the effects of CAF CM supplementation(Fig. 3D). ASNase treatment significantly inhibited tumor cell proliferation, and this inhibitory effect was not reversed by CAF CM supplementation. Moreover, ASNase completely abolished the ability of CAF CM to rescue CTO-induced growth inhibition. Notably, the combination of ASNase and CTO further suppressed tumor cell proliferation compared with either treatment alone. Western blot analysis showed that degradation of ASN in CAF CM led to stronger activation of ISR signaling under CTO stress, indicating that tumor cells experienced greater metabolic stress(Fig. 3 E-G). Interestingly, the combination of CTO and ASNase led to significant increase in p-eIF2α expression, however, its downstream ATF4 expression remained suppressed. Meanwhile, c-Myc expression was further reduced under the combined treatment condition. These results suggests that global translation remain suppressed, however, severe metabolic stress impaired the ATF4-mediated adaptive program under combination treatment. Together, these findings suggest that CAFs support tumor cell survival under OXPHOS inhibition by supplying ASN, which both alleviates ISR activation and support ISR downstream effects.

Additionally, intracellular metabolomic analysis was performed comparing untreated tumor cells, CAF CM-treated tumor cells, and CAF CM + ASNase-treated tumor cells(Fig. S1A-E). Interestingly, degradation of ASN significantly altered the intracellular metabolic landscape even when other metabolites remained abundant. Several lipid metabolites were upregulated(Fig. S1C), whereas short-peptide synthesis was markedly suppressed(Fig. S1E). These findings suggest that depletion of ASN significantly reshapes intracellular metabolic pathways even in the presence of other extracellular metabolites.

### 4. ASNase sensitizes PDAC to OXPHOS inhibition therapy *in vivo*

To determine whether ASN also contributes to resistance to OXPHOS inhibition therapy *in vivo*, we evaluated the therapeutic effect of combining CTO with Oncaspar (PEG-L-asparaginase)(Fig. 4A-B). The results showed that the combination treatment significantly reduced tumor weight and inhibited tumor growth compared with either treatment alone. Immunohistochemical analysis further demonstrated that Ki67 expression in tumor tissues was markedly decreased in the combination treatment group, indicating strong anti-proliferative activity(Fig. 4C-D).

**Figure 4.**
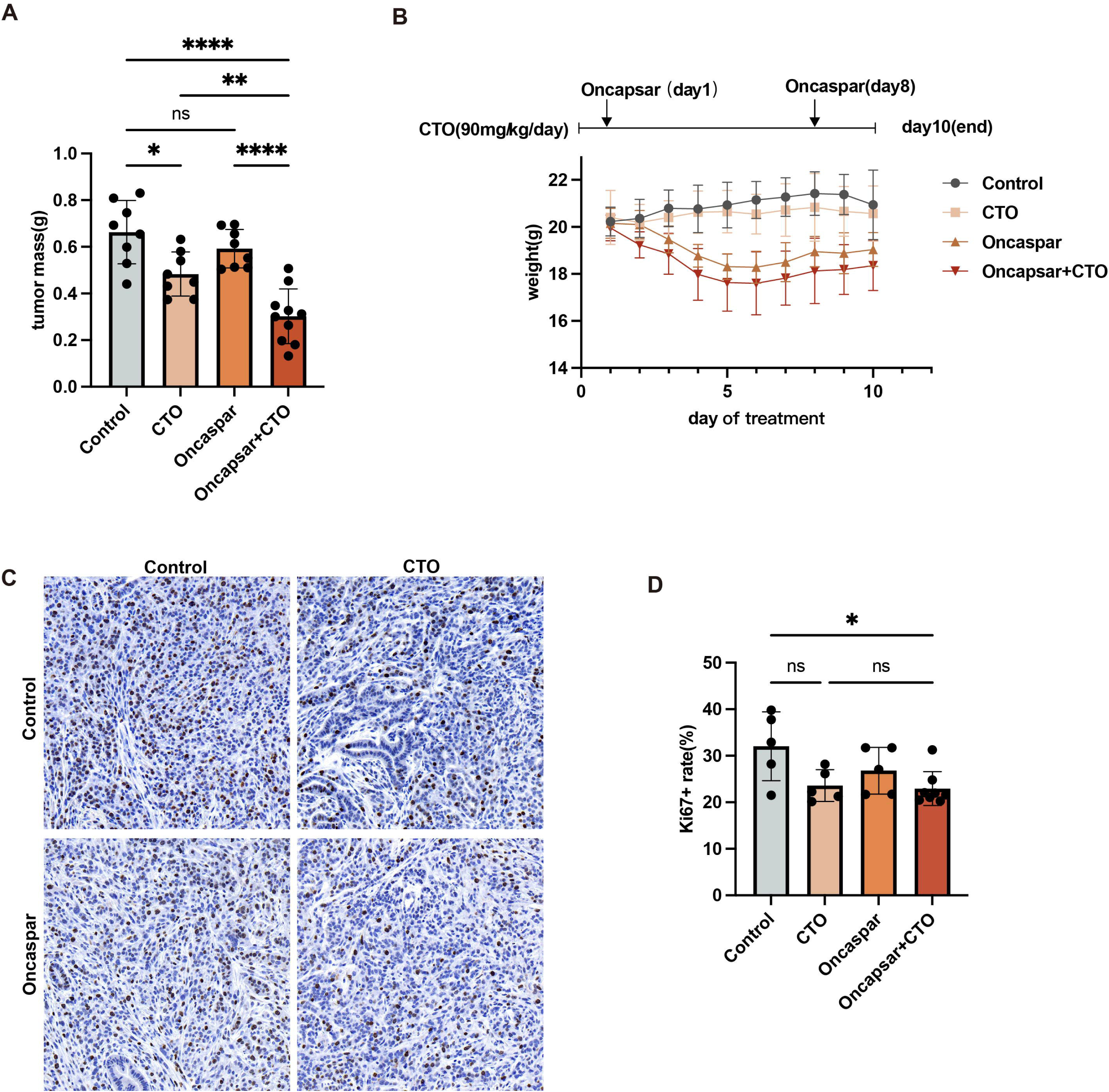
Combination treatment suppresses tumor growth in an orthotopic pancreatic cancer mouse model. A. Tumor weights from the orthotopic KPC pancreatic cancer model in C57 mice. B. Experimental design showing drug administration schedule and body weight changes of mice during treatment. C. Representative Ki67 immunohistochemical staining images of tumor sections. D. Quantification of Ki67-positive cells in tumor sections. Control (n = 5), CTO (n = 5), Oncaspar (n = 5), Oncaspar + CTO (n = 8) (t test, p < 0.05).

### 5. SLC38A4 functions as a key amino acid transporter under OXPHOS stress in PDAC

To identify more specific therapeutic targets, we further analyzed transcriptomic data from tumor cells treated with CTO with or without CAF CM (Fig. 5A). Among genes associated with ASN metabolism, the sodium-dependent neutral amino acid transporter *Slc38a4* was significantly upregulated in response to CAF CM treatment. Further analysis confirmed that CAF CM significantly increased *Slc38a4* expression at both the mRNA and protein levels(Fig 5. B). Notably, the increase in SLC38A4 expression closely correlated with c-Myc activation(Fig. 5C). Previous ChIP-seq studies have reported that c-Myc can function as an upstream transcription factor of SLC38A4 in several cell types, although this regulatory relationship has not yet been reported in pancreatic cancer cells. Therefore, we hypothesize c-Myc may regulate *Slc38a4* expression.

**Figure 5.**
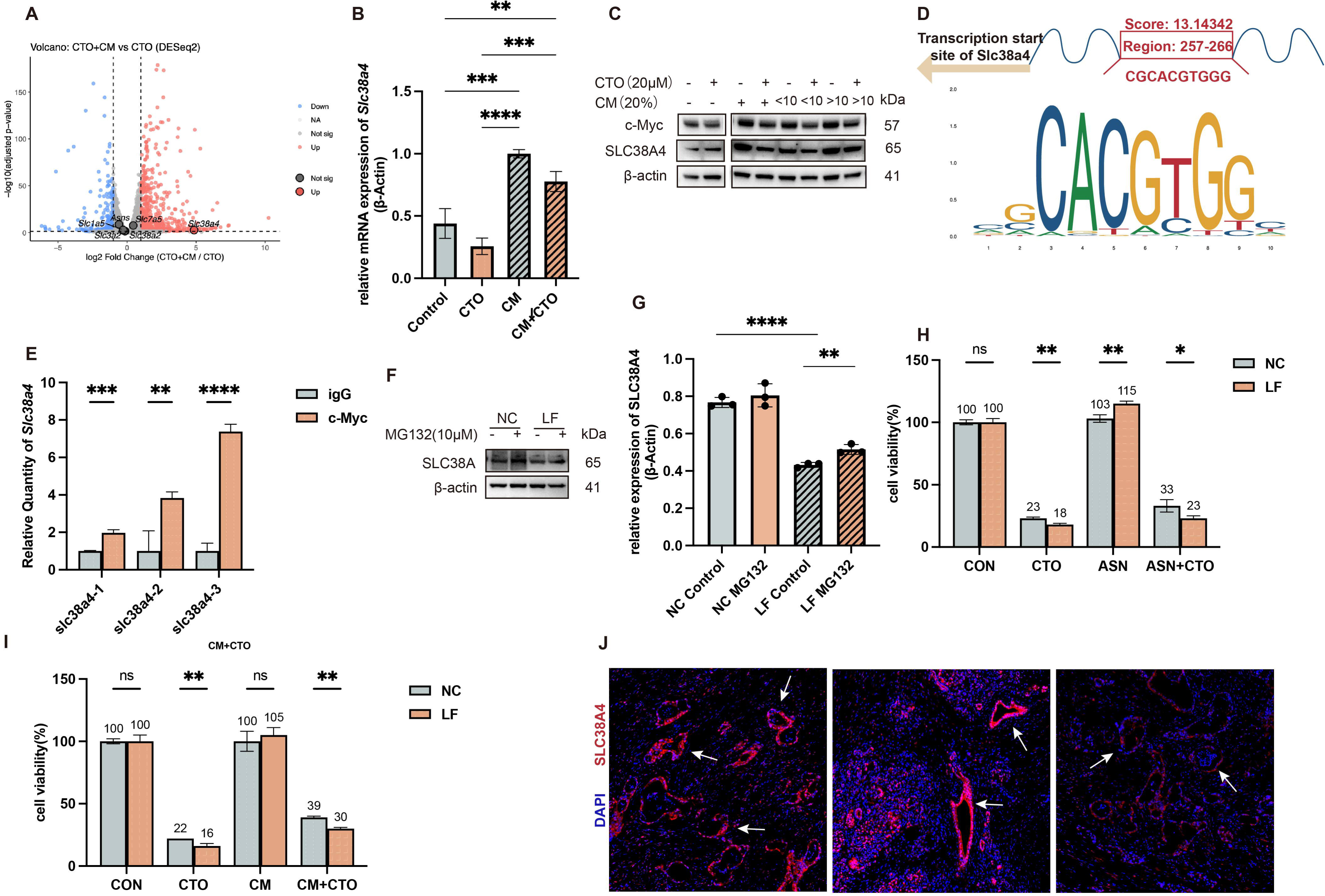
CAF-induced c-Myc upregulates SLC38A4 to support metabolic adaptation. A. Transcriptomic analysis of Pan02 cells. Volcano plot showing differential gene expression between CTO and CTO + CAF conditioned medium groups, with Slc38a4 significantly upregulated. B. qPCR validation in Pan02 cells showing increased Slc38a4 mRNA expression following CAF conditioned medium treatment (t test, n = 3, ****p < 0.0001, ***p < 0.001, **p < 0.01). C. Representative Western blot images showing c-Myc and SLC38A4 expression in Pan02 cells. D. Motif prediction of c-Myc binding sites in the *Slc38a4* promoter region using the JASPAR database. E. CUT&Tag-qPCR results validating c-Myc binding to the *Slc38a4* promoter (t test, n = 3, ****p < 0.0001, ***p < 0.001, **p < 0.01). F. Western blot showing SLC38A4 expression following gene knockdown in KPC cells. G. Quantification of Western blot results shown in panel F using ImageJ (t test, n = 3, ****p < 0.0001, **p < 0.01). H–I. SRB proliferation assays showing the effect of SLC38A4 knockdown on KPC cell proliferation (t test, n = 3, **p < 0.01, *p < 0.05). J. Immunofluorescence staining of SLC38A4 in pancreatic cancer patient samples (SLC38A4, red; DAPI, blue).

To investigate this possibility, potential c-Myc binding sites within the *Slc38a4* promoter region were predicted using the JASPAR database(Fig. 5D). CUT&Tag–qPCR analysis demonstrated significant enrichment of c-Myc binding at the *Slc38a4* promoter compared with the IgG control, indicating direct transcriptional regulation(Fig. 5E).

To further evaluate the functional role of SLC38A4, we attempted to generate an overexpression model by lentiviral infection. Although SLC38A4 mRNA levels increased markedly following transduction, the protein level was unexpectedly reduced. Treatment with the proteasome inhibitor MG132 for 4 hours restored SLC38A4 protein abundance, suggesting that the ectopically expressed protein undergoes proteasome-mediated degradation, possibly due to protein misfolding (Fig. 5F-G). Therefore, this system effectively generated a low-function SLC38A4 model for subsequent functional analyses. Cells with reduced SLC38A4 function exhibited increased sensitivity to CTO treatment. Moreover, the ability of ASN supplementation or CAF CM to rescue CTO-induced growth inhibition was markedly reduced in this model.(Fig. 5H-I)

These findings demonstrate that SLC38A4 plays a critical role in tumor cell adaptation to OXPHOS stress in PDAC(Fig. 5J). Furthermore, analysis of human clinical samples confirmed that SLC38A4 is specifically expressed in pancreatic ductal epithelial cells, suggesting its potential as a tumor-selective therapeutic target.

## Discussion

In this study, we investigated metabolic interactions between CAFs and PDAC cells under pharmacological inhibition of oxidative phosphorylation. Our findings demonstrate that PDAC cells activate ISR when mitochondrial respiration is suppressed. However, metabolites supplied by CAFs can partially relieve this metabolic stress, thereby supporting tumor cell survival and proliferation. Through metabolomic analyses, we identified ASN as a key metabolite that supports tumor cell adaptation to mitochondrial metabolic stress. Notably, combined treatment with ASNase and CTO revealed that the execution of the ISR downstream adaptive program requires a minimal supply of extracellular ASN. In the absence of ASN, although eIF2α phosphorylation and global translation attenuation were maintained, the ATF4-dependent adaptive transcriptional program collapsed, leading to simultaneous depletion of both intracellular and extracellular metabolic support. These findings suggest that ISR activation alone does not necessarily promote tumor survival. Instead, the outcome of ISR signaling may depend on the availability of key metabolic substrates such as ASN. When extracellular ASN supply is limited, the ATF4-dependent adaptive program fails, converting ISR from a pro-survival pathway into a vulnerability. Furthermore, we identified the amino acid transporter SLC38A4 as a critical mediator facilitating this tumor-stromal metabolic coupling. Collectively, these findings highlight a previously underappreciated role of amino acid transport and stromal metabolic support in enabling PDAC cells to adapt to metabolic therapies.

Amino acid availability in cells is primarily monitored by two major signaling systems with opposing responses: the GCN2 pathway and the mechanistic target of rapamycin complex 1 (mTORC1). GCN2 is activated under amino acid deprivation, whereas mTORC1 activity is suppressed under the same conditions. In PDAC, previous studies have reported that tumor cells can activate the GCN2-eIF2α-ATF4 signaling axis to upregulate asparagine synthetase (ASNS), thereby replenishing intracellular ASN levels and partially restoring mTORC1 activity during respiratory stress (Pathria et al., 2019; Krall et al., 2021). Our findings are consistent with the importance of this pathway in enabling tumor cells to tolerate OXPHOS inhibition. However, our study further suggests that extracellular ASN supply may represent an additional and potentially critical layer of regulation. Compared with endogenous ASNS-mediated synthesis, ASN derived from the tumor microenvironment may provide a more immediate metabolic resource that allows tumor cells to overcome metabolic stress. Although our experiments did not directly quantify the exact contribution of CAFs to total ASN levels within TME, CAFs may represent an important source of extracellular ASN within the TME. In contrast, aspartate, which exists in dynamic equilibrium with ASN in intracellular metabolism, plays an important role in tumor metabolism but may primarily maintained through endogenous metabolic processes rather than extracellular supply (Sullivan et al., 2018). Our results further suggest that the ATF4-ASNS axis requires a minimal level of ASN, which may function both as a signaling metabolite and as a precursor supporting aspartate-related metabolic pathways. Together with our findings, these observations suggest that the GCN2-eIF2α pathway in PDAC may be strongly influenced not only by intrinsic metabolic activity but also by nutrient availability within TME, highlighting the importance of stromal nutrient supply in shaping metabolic stress responses in PDAC.

Beyond the effect of a single metabolite, our metabolomic analyses suggest that ASN depletion may trigger broader alterations in intracellular metabolic networks(Fig S1). Many previous studies in tumor metabolism have focused on the function of individual metabolites; however, our findings emphasize the dynamic balance of metabolic networks within cancer cells. Changes in the availability of a single metabolite can reshape multiple interconnected metabolic pathways. Despite increasing interest in tumor metabolism, how metabolite exchange within TME reshapes intracellular metabolic networks remains poorly understood. From a translational perspective, strategies aimed at depriving tumor cells of ASN, such as the use of ASNase, have shown promising potential in combination with metabolic inhibitors, including OXPHOS inhibitors, particularly those targeting mitochondrial Complex I(Krall et al. 2021). In our study, the combination of ASNase and CTO significantly suppressed tumor growth. However, we also observed a severe reduction in body weight in treated mice. Although this weight loss did not affect overall activity during the experimental period and remained within the predefined humane endpoints approved for the animal study, it nevertheless raises potential safety concerns. Given that many PDAC patients already suffer from cancer-associated cachexia, systemic ASN depletion may carry clinical risks. Therefore, identifying more selective targets that disrupt ASN-dependent metabolic adaptation without inducing severe systemic toxicity represents an important future direction.

In our study, SLC38A4 was identified as a potential key mediator by screening genes associated with ASN metabolism. However, it should be noted that SLC38A4 is not a specific transporter for ASN but rather a sodium-dependent transporter for a range of neutral amino acids. Therefore, although our data suggest that changes in SLC38A4 expression sensitize tumor cells to OXPHOS inhibition, it remains unclear whether this effect is entirely dependent on ASN transport. Indeed, several metabolites transported by SLC38A4 may also contribute to tumor metabolism. For example, alanine, one of the substrates of SLC38A4, has been reported to support tumor cell metabolism and proliferation in previous research in PDAC through stromal-derived nutrient exchange. Thus, the impact of SLC38A4 on tumor cell growth and drug resistance may involve additional metabolic mechanisms beyond ASN transport, which require further investigation. Notably, the function of SLC38A4 appears to exhibit strong tissue specificity and context dependence. Previous studies have reported that the expression and functional consequences of SLC38A4 differ across tumor types, including hepatocellular carcinoma and colorectal cancer. Such discrepancies may be explained by organ-specific metabolic and stress-response environments. The pancreas is characterized by extremely high rates of protein synthesis and secretion, particularly in acinar and endocrine cells, making it especially sensitive to disruptions in amino acid homeostasis and ISR signaling. Moreover, pancreatic injury and regeneration processes, including acinar-ductal metaplasia (ADM), autophagy activation, and dynamic regulation of mTORC1 signaling, are tightly linked to amino acid sensing and stress adaptation(Grippo and Sandgren 2012; Wei et al. 2016; Willet et al. 2018). These biological features suggest that nutrient sensing and amino acid transport may play particularly important roles in pancreatic tissue. Consequently, in PDAC, the function of SLC38A4 may extend beyond basic amino acid transport and instead participate more deeply in metabolic adaptation and tumor-stromal symbiosis under therapeutic stress. In contrast, tissues such as the liver possess greater metabolic redundancy and distinct oncogenic drivers, which may lead to different functional outcomes for SLC38A4 in hepatic tumors. This context-dependent role may help explain the heterogeneous observations reported in other cancers, such as colorectal cancer. It also raises the possibility that SLC38A4 may play distinct roles in primary pancreatic tumors and metastatic lesions, particularly in the liver. Therefore, further *in vivo* studies will be required to clarify the functional significance of SLC38A4 before therapeutic strategies targeting this transporter can be considered for PDAC treatment.

Nevertheless, several limitations of this study should be acknowledged. First, although our work focused primarily on ASN supplied by CAFs, ASN within TME is unlikely to originate exclusively from stromal fibroblasts. Systemic sources, including dietary ASN and circulating amino acids, have also been reported to contribute to tumor metabolism in previous studies. Therefore, the relative contribution of CAF-derived ASN compared with systemic ASN supply remains to be clarified. Second, our study mainly investigated the metabolic interaction between CAFs and tumor cells and did not explore the potential impact of ASN metabolism on the tumor immune microenvironment. Given that amino acid availability can influence immune cell function, it will be important for future studies to determine whether ASN depletion or SLC38A4-mediated transport also affects anti-tumor immunity. Third, although we identified SLC38A4 as a potential mediator of tumor adaptation to OXPHOS inhibition, the upstream regulatory mechanisms controlling SLC38A4 expression in PDAC remain incompletely understood. In addition, it remains unclear whether other amino acid transporters participate in similar metabolic adaptations. Future studies integrating metabolic flux analysis, genetic knockout models, and clinical sample validation will be required to clarify the precise role of SLC38A4 in PDAC metabolic adaptation. If SLC38A4 is confirmed to be a key mediator enabling tumor cells to escape ISR-induced metabolic stress, targeting SLC38A4 in combination with OXPHOS inhibitors or chemotherapy may represent a promising strategy to disrupt tumor-stromal metabolic symbiosis and overcome therapeutic resistance in PDAC.

## Author Contributions

Ziyi Qin and Shengxi Li contributed equally to this work. Ziyi Qin performed the majority of the experiments, conducted data analysis, generated the figures, and wrote the manuscript。 Shengxi Li participated in experimental execution. Yeting Xu contributed to experimental design, performed experiments, and assisted with data analysis. Jiaming Zou, Jinyang Ma, Yingying Wang, and Yucheng Wang participated in experimental work. Rui Ju, Lin Wang, and Lei Guo conceived and supervised the study, and provided critical revisions of the manuscript.

## Data Availability

The datasets generated during and/or analysed during the current study are available from the corresponding author on reasonable request.

---

**Supplementary Figure 1.** Metabolomic profiling of KPC cells. A. Heatmap showing enrichment of TCA cycle–related metabolites in Pan02 cells based on metabolomic analysis. B. Heatmap showing enrichment of amino acid–related metabolites in Pan02 cells. C. Heatmap showing enrichment of lipid-related metabolites in Pan02 cells. D. Heatmap showing enrichment of nucleotide-related metabolites in Pan02 cells. E. Heatmap showing enrichment of short peptide–related metabolites in Pan02 cells.

