## Supplementary figures and images for "Stromal asparagine supports tumor adaptation to oxidative phosphorylation inhibition through SLC38A4-mediated metabolic coupling"

### Supplementary Figure1

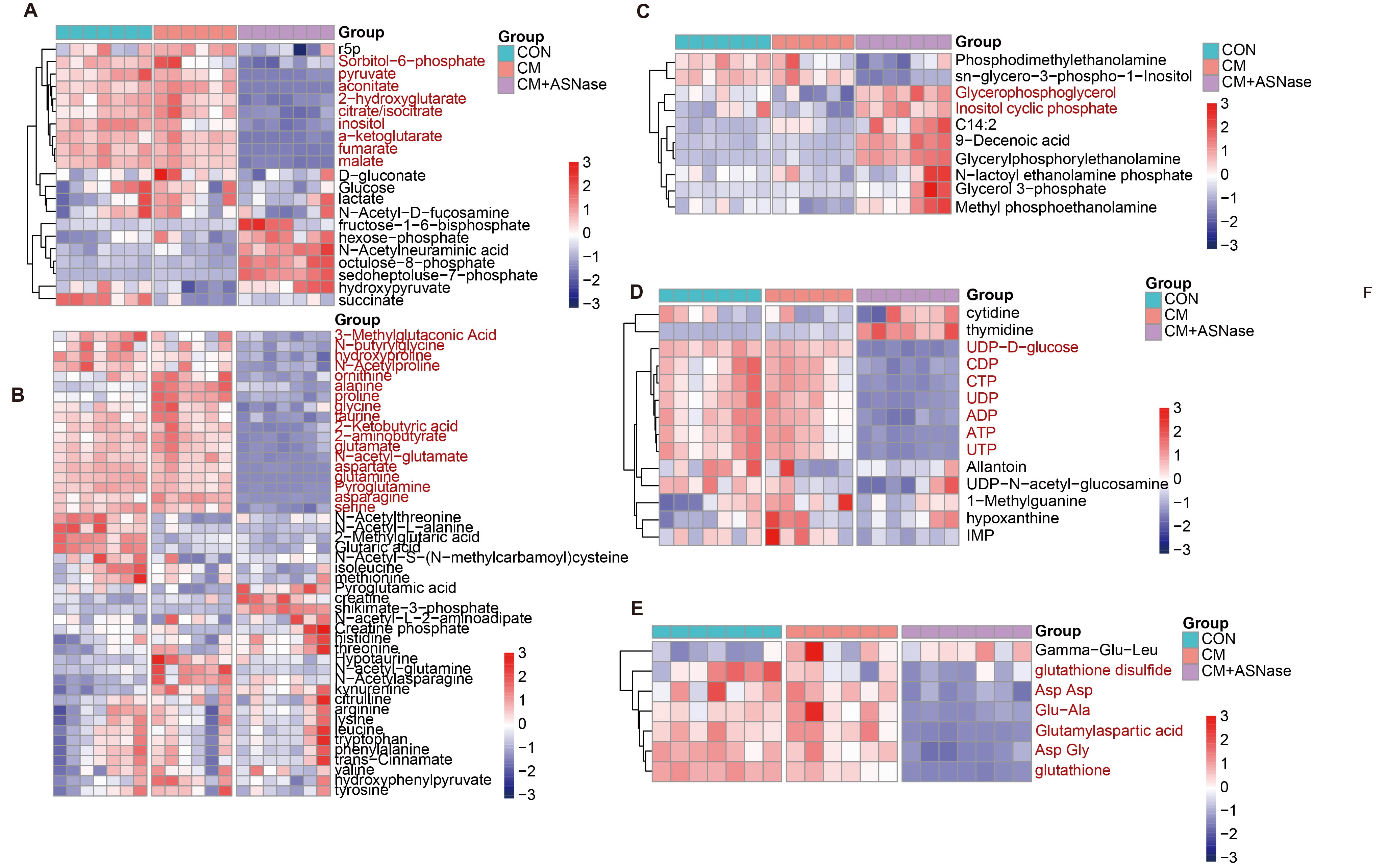
