## Supplementary Table1 for "Stromal asparagine supports tumor adaptation to oxidative phosphorylation inhibition through SLC38A4-mediated metabolic coupling"

| Metabolites meeting all screening criteria | Metabolites meeting partial screening criteria |
| --- | --- |
| asparagine | O-phosphoethanolamine |
| succinate | alanine |
| proline | aspartate |
| muramic acid | phospho-dimethylethanolamine |
| | histidine |
| | AMP |
| | GMP |
| | phosphate |
| | phosphorylcholine |
| | PC(14:0/18:2) |
| | PE(18:2/18:1) |
